# TMEM106B-mediated SARS-CoV-2 infection allows for robust ACE2-independent infection *in vitro* but not *in vivo*

**DOI:** 10.1101/2024.05.08.593110

**Authors:** Kexin Yan, Troy Dumenil, Romal Stewart, Cameron R. Bishop, Bing Tang, Andreas Suhrbier, Daniel J. Rawle

## Abstract

Angiotensin converting enzyme 2 (ACE2) serves as the primary entry receptor for severe acute respiratory syndrome coronavirus 2 (SARS-CoV-2). However, ACE2-independent entry has been observed *in vitro* for SARS-CoV-2 strains containing the E484D amino acid substitution in the spike protein. In this study, we conducted a whole genome CRISPR-Cas9 knockout screen using a SARS-CoV-2 strain containing the spike-E484D substitution (SARS-CoV-2_MA1_) to identify the ACE2-independent entry mechanisms. Our findings revealed that SARS-CoV-2_MA1_ infection in HEK293T cells relied on heparan sulfate and endocytic pathways, with TMEM106B emerging as the most significant contributor. While SARS-CoV-2_MA1_ productively infected human brain organoids and K18-hACE2 mouse brains, it did not infect C57BL/6J or *Ifnar^-/-^* mouse brains. This suggests that ACE2-independent entry via TMEM106B, which is a protein that is predominantly expressed in the brain, did not overtly increase the risk of SARS-CoV-2 neuroinvasiveness in wild-type mice. Importantly, SARS-CoV-2_MA1_ did not replicate in *Ace2^-/-^* mouse respiratory tracts. Overall, this suggests that robust ACE2-independent infection by SARS-CoV-2_E484D_ is likely a phenomenon specific to *in vitro* conditions, with no apparent clinical implications.

## INTRODUCTION

Angiotensin converting enzyme 2 (ACE2) is the key entry receptor for severe acute respiratory syndrome coronavirus 2 (SARS-CoV-2)^1^. In an infected cell, furin cleaves spike (S) at the S1/S2 boundary, with S1 and S2 remaining non-covalently linked and assembled on the surface of virions^2^. The SARS-CoV-2 spike receptor binding domain (RBD) binds to the ACE2 receptor, with Transmembrane Serine Protease 2 (TMPRSS2) then able to cleave S2’ to expose the fusion peptide to trigger viral fusion with the host cell membrane^3^. Pre-cleavage of S1/S2 by furin is required for efficient TMPRSS2 cleavage of S2’, and SARS-CoV-2 with deletions or changes in the S furin cleavage site (FCS) favor entry via endosomes where S2’ is instead cleaved by cathepsin L for fusion with the endosomal membrane^3^. Furin cleavage has been shown as important for transmission between ferrets^3^, suggesting that furin cleavage and TMPRSS2-mediated fusion at the cell membrane are preferred over endosomal entry in respiratory tract cells. *In vitro* experiments have proposed a series of alternative or additional receptors for SARS-CoV-2 entry^4^, including DC-SIGN^5^, AXL^6^, neuropilin-1^7^, and CD147^8^. However the ability for robust ACE2-independent infection has not been demonstrated in an *in vivo* respiratory tract.

We previously identified a SARS-CoV-2 strain, which we called SARS-CoV-2 mouse adapted 1 (SARS-CoV-2_MA1_), that was capable of robust infection in ACE2^-/-^ cell lines *in vitro*^9^. We and others have identified that the E484D substitution in the spike protein was key for robust ACE2-independent infection^9–13^. Spike D484 has been identified in SARS-CoV-2 sequences from 337 human samples in the GISAID database (epicov.org), which is just ∼0.002% of all samples and suggests no overt selective advantage in humans. SARS-CoV-2_MA1_ also contains the QTQTN deletion in the spike FCS^9^, which impairs furin and TMPRSS2 cleavage^14^, suggesting infection is primarily via the endosomal entry route after engaging a non-ACE2 receptor. While deletions in the spike FCS often occurs during passage in TMPRSS2-negative cells *in vitro*^15^, deletions in the FCS are not commonly identified in human samples^9^. SARS-CoV-2_MA1_ could infect a range of ACE2-negative cell lines^9^ (HEK293T, 3T3, AE17, BHK-21, A549, HeLa, LLC-PK1, and Caco2-ACE2^-/-^), although the mechanism was unknown, and whether ACE2-independent infection occurred *in vivo* was not determined.

Herein we sought to identify the mechanisms of ACE2-independent infection by SARS-CoV-2_MA1_, and determine the relevance of this entry mechanism *in vivo*. We identified heparan sulfate biosynthesis and endosomal pathways as the cellular pathways involved in SARS-CoV-2_MA1_ entry in the absence of ACE2, with TMEM106B the main contributor. During the course of our study, Baggen *et al* also identified TMEM106B as an alternative receptor engaged by spike containing the E484D substitution^13^. However, they did not demonstrate that this entry mechanism was functional *in vivo*. Herein we show that SARS-CoV-2_MA1_ cannot productively infect the respiratory tract in *Ace2^-/-^* mice. Overall, our study identifies and confirms TMEM106B as an alternative SARS-CoV-2_E484D_ receptor *in vitro*, but not in mouse respiratory tracts.

## RESULTS

### Whole genome CRISPR-Cas9 knockout screening identified TMEM106B as a major facilitator of ACE2-independent infection

To identify the key host factors that facilitate ACE2-independent infection for SARS-CoV-2_MA1_, we performed a whole genome CRISPR-Cas9 knockout screen in HEK293T cells. HEK293T cells express little or no ACE2 or TMPRSS2 and do not support infection of ACE2-dependent SARS-CoV-2 isolates^9^. HEK293T cells were transduced with the Brunello human CRISPR Knockout Pooled Library^16^, and were then infected with SARS-CoV-2_MA1_ (Fig. 1A). Infection caused near-complete cytopathic effect by day 3 post infection, with clusters of surviving cells visible by day 11 post infection. Surviving cells were expanded and re-infected with SARS-CoV-2_MA1_, and complete resistance to infection was indicated by the lack of any cytopathic effect. DNA from surviving cells were extracted, and barcoded sgRNAs were amplified by PCR and identified by sequencing using the NextSeq 2000 platform. Seven sgRNAs were significantly enriched in the surviving cells; TMEM106B, SLC35B2, CCZ1B, EXT2, CCZ1, B4GALT7, and EXT1 (Fig. 1B). Of these, SLC35B2, B4GALT7, EXT1 and EXT2 are involved in heparan sulfate biosynthesis pathways^17^, and TMEM106B, CCZ1, and CCZ1B are involved in endocytic pathways^18,19^. These host factors have also been identified in other SARS-CoV-2 CRISPR screening experiments^20–25^. Our data now implicates the role of heparan sulfate attachment and endosomal entry for SARS-CoV-2 infection in an ACE2 and TMPRSS2 negative setting.

**Figure 1.**
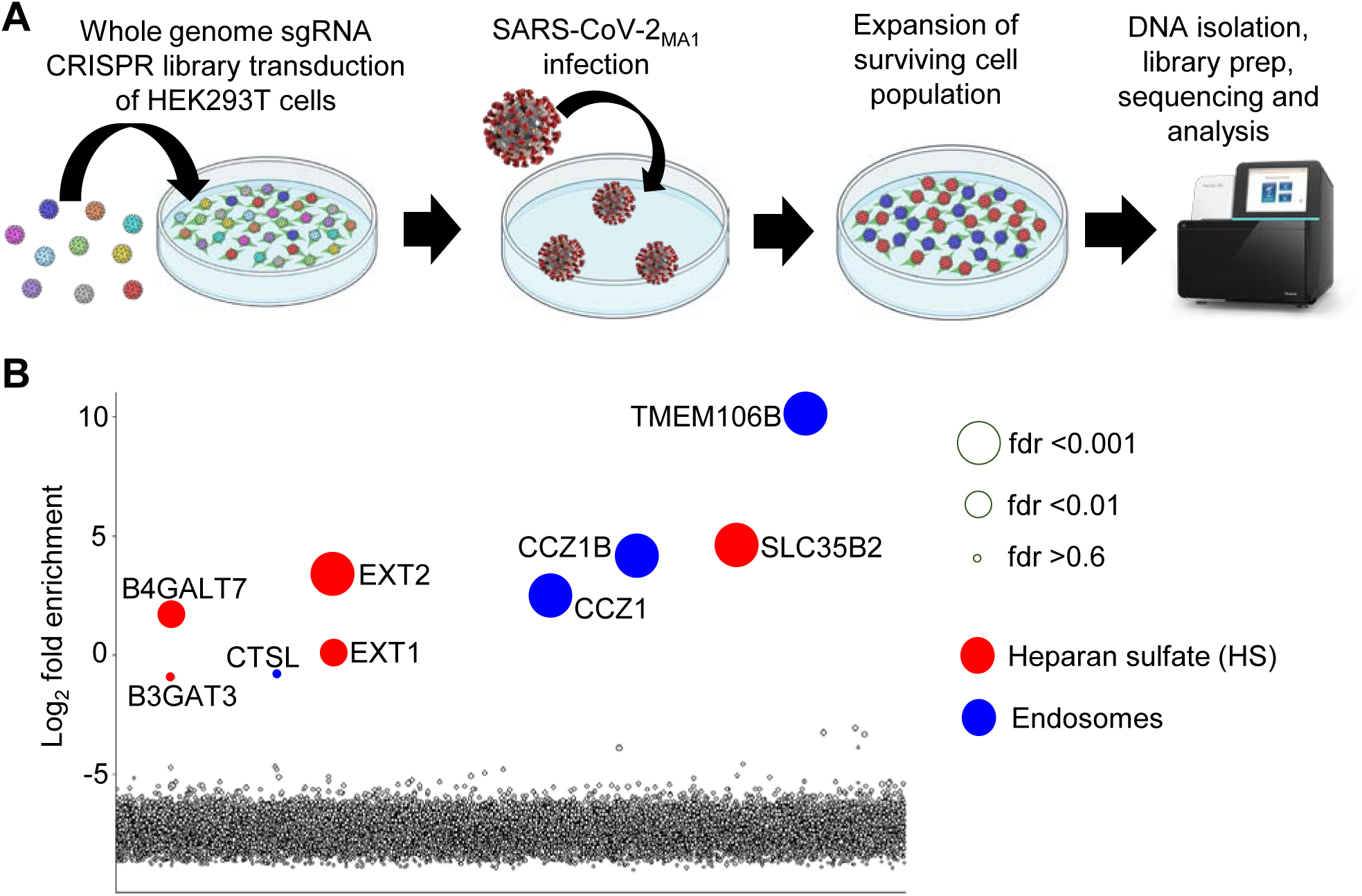
Whole genome CRISPR-Cas9 knockout screen identifies heparan sulfate and endosomal genes as facilitators of ACE2-independent SARS-CoV-2 infection. A) Overview of steps in whole genome CRISPR-Cas9 knockout screening in HEK293T cells to identify genes that facilitate ACE2-independent infection by SARS-CoV-2_MA1_. B) Data from whole genome CRISPR-Cas9 knockout screen in HEK293T cells. Each circle represents a gene, with size scaled for false discovery rate (fdr) as indicated. Circles colored red indicate genes involved in heparan sulfate biosynthesis pathways, and blue indicates genes involved in endosomal pathways. Genes are distributed randomly across the x-axis. The y-axis represents the log_2_ fold enrichment comparing post-selection (21 days post SARS-CoV-2_MA1_ infection) with pre-selection (prior to infection). Genes are labelled if the log_2_ fold enrichment was higher than -3. Data is the average of three sequencing reactions for pre-infection (pooled from five independent biological replicates), and four sequencing reactions for post-infection (from four independent biological replicates), calculated using Robust Rank Aggregation algorithm in MAGeCK software.

The most significantly enriched sgRNA was TMEM106B, which had a log_2_ fold change more than twice the next highest gene (Fig. 1B). During the course of our study, Baggen *et al* identified that their CRISPR-Cas9 knockout screen which identified TMEM106B as an important pro-viral factor for SARS-CoV-2^22^ was unknowingly performed using a SARS-CoV-2 strain that acquired the E484D mutation during passage in Huh7 cells^13^. They showed that TMEM106B binds directly to the SARS-CoV-2_E484D_ receptor binding domain (RBD) to facilitate entry in the absence of ACE2^13^. This is consistent with our findings (Fig. 1B), and implicates TMEM106B in the ACE2-independent entry of SARS-CoV-2_MA1_^9^.

### SARS-CoV-2_MA1_ efficiently infects human cortical brain organoids and RENcell VM neural progenitor cells

Since TMEM106B is primarily expressed in brain tissue (www.proteinatlas.org) and involved in neurodegenerative disorders^19,26–29^, we sought to further investigate SARS-CoV-2_MA1_ replication in neural cells. Firstly, we infected 2-3 mm diameter human cortical brain organoids (hBOs) generated from human induced pluripotent cells (hiPSCs) in a rotating incubator^30^. SARS-CoV-2_MA1_ replicated to high titers in hBOs (Fig. 2A), and caused robust cytopathic effect by day 9 and 12 post infection (Fig. 2B). SARS-CoV-2_MA1_ reached ∼5.5 log_10_ CCID_50_/ml in the supernatant by day 2 post infection, which was significantly higher than its parental SARS-CoV-2_QLD02_ strain (Fig. 2A) and the Omicron BA.1, BA.5, and XBB variants^30^. Immunohistochemistry of infected hBOs using an anti-spike monoclonal antibody^31^ showed that SARS-CoV-2_MA1_ infected cells were primarily localized to the outer surface of the organoids (Fig. 2C), where the cells are in direct contact with the culture medium. Hematoxylin and eosin staining showed loss of integrity of the outer membrane of the hBOs (Fig. 2C). SARS-CoV-2_MA1_ also replicated to high titers in 2D cultures of RENcell VM neural progenitor cells and caused robust cytopathic effect (Supplementary Figure 1).

**Figure 2.**
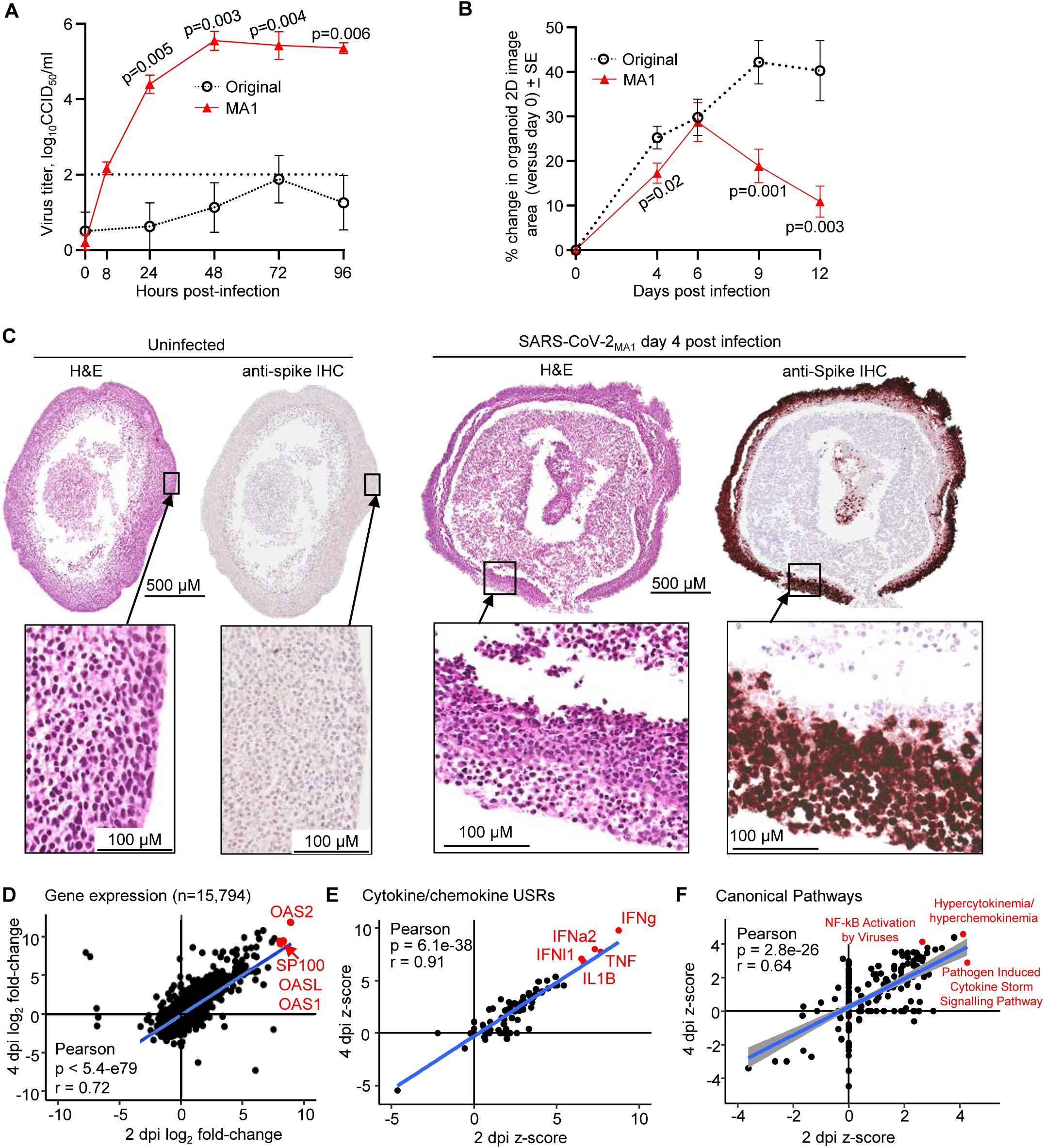
SARS-CoV-2_MA1_ efficiently infects human cortical brain organoids. A) Viral titers in the supernatant of brain organoid cultures sampled at the indicated hours after infection with original SARS-CoV-2_QLD02_ (black circles) or SARS-CoV-2_MA1_ (red triangles). Data represents the mean (MA1; n=10 at 0, 24, and 48 hpi, n=7 at 72 and 96 hpi, and n=3 at 8 hpi. Original; n=4 at all time points), and error bars represent standard error. B) Mean percentage change from day 0 in organoid area (calculated by ImageJ as the area within the circumference of the 2D image) at days 4, 6, 9 and 12 post infection. Data is the mean of 4 organoids per group and error bars represent standard error. C) Hematoxylin and eosin (H&E) and anti-spike immunohistochemistry (IHC) of brain organoids, either uninfected (left) or at day 4 post SARS-CoV-2_MA1_ infection (right). Dark brown stain in IHC images indicates cells expressing SARS-CoV-2_MA1_ spike protein. Images are representative of at least 4 brain organoids. D) The log_2_ fold change of all genes (n=15,794) for SARS-CoV-2_MA1_ infected organoids compared to uninfected organoids is shown for day 2 (x-axis) versus day 4 (y-axis). Statistics by Pearson correlation. E) Ingenuity Pathway Analysis (IPA) cytokine or chemokine Up-Stream Regulators (USRs) z-scores are shown for day 2 (x-axis) versus day 4 (y-axis). Statistics by Pearson correlation. F) IPA Canonical Pathways z-scores are shown for day 2 (x-axis) versus day 4 (y-axis). Statistics by Pearson correlation.

### SARS-CoV-2_MA1_ replication in human brain organoids triggers an innate cytokine storm-like transcriptional response

To determine transcriptional responses to SARS-CoV-2_MA1_ infection in hBOs, we performed RNAseq at day 2 and 4 post infection and performed differential expression analysis compared to uninfected hBOs. SARS-CoV-2_MA1_-induced changes in the transcriptome were significantly correlated for day 2 and 4 post-infection, with interferon stimulated genes dominating the most highly upregulated genes (Fig. 2D, Supplementary Table 2). Using an fdr threshold of <0.05, there were 9126 differentially expressed genes (DEGs) on day 2 and 11211 on day 4, indicating a strong transcriptional response to SARS-CoV-2_MA1_ infection in hBOs. To generate smaller DEG lists suitable for pathway analysis, we included a log_2_ fold change threshold of >1 or <-1 in addition to fdr <0.05. Using this criteria, there were 1115 DEGs on day 2 and 2986 DEGs on day 4 (Supplementary Table 2), which were analysed using Ingenuity Pathway Analysis (IPA). The cytokine/chemokine Up-Stream regulators (USRs) significantly correlated on day 2 and 4 post infection and were dominated by interferons and pro-inflammatory cytokines (Fig. 2E). Similarly, Canonical Pathways analyses indicated a significant upregulation in cytokine storm-like transcriptional responses on day 2 and 4 post infection (Fig. 2F). Overall, SARS-CoV-2_MA1_ infected hBOs significantly higher than ACE2-dependent SARS-CoV-2, and this triggered a strong interferon and pro-inflammatory cytokine response.

### SARS-CoV-2_MA1_ has increased replication in K18-hACE2 mouse brains, but not C57BL/6J or *Ifnar^-/-^* mouse brains

Since SARS-CoV-2_MA1_ had productive infection of human cortical brain organoids, and the spike E484D substitution in SARS-CoV-2_MA1_ engages the brain-expressing TMEM106B receptor^13^ (Fig. 1B), we sought to determine whether SARS-CoV-2_MA1_ had increased propensity to infect brains *in vivo*. K18-hACE2 mice express the human ACE2 receptor from the K18 promoter, which was an important mouse model for original SARS-CoV-2 as the spike of this virus does not engage mouse ACE2^32^. K18-hACE2 mice allow for SARS-CoV-2 to enter the brain via infection of the olfactory epithelium^33,34^, often leading to fulminant brain infection^30^.

K18-hACE2 mice were given an intrapulmonary inoculum via the intranasal route of SARS-CoV-2_QLD02_ or SARS-CoV-2_MA1_. Infection led to a rapid decline in body weight (Fig. 3A) and increase in disease scores (Fig. 3B), with almost all mice requiring humane euthanasia on day 4 post infection (Fig. 3C). Viral titers in lungs, nasal turbinates and brains were measured by CCID_50_ assay on day 4 post infection. MA1 had significantly higher viral titers in lungs (∼0.7 log higher) and brains (∼1.8 logs higher), while viral titers in nasal turbinates were not significantly different (Fig. 3D).

**Figure 3.**
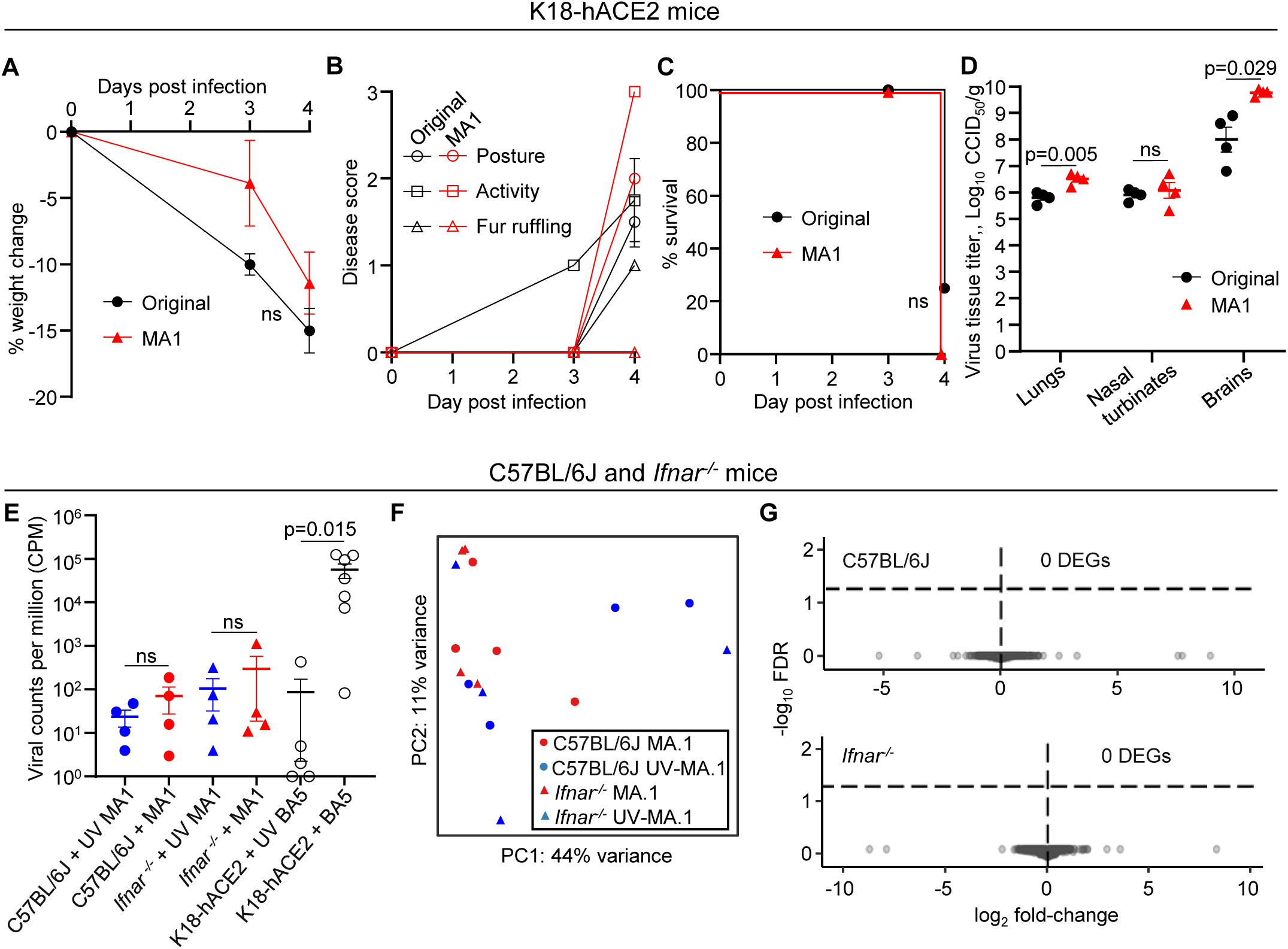
SARS-CoV-2_MA1_ has no overt increased risk of neurotropism in C57BL/6J or *Ifnar^-/-^* mice. A) Percent weight change from day 0 after intranasal infection of K18-hACE2 mice with original SARS-CoV-2_QLD02_ (black circles) or SARS-CoV-2_MA1_ (red triangles). Data is the mean of 4 mice per group and error bars represent standard error. B) Clinical disease scores for posture (circles), activity (squares) and fur ruffling (triangles) were monitored in K18-hACE2 mice on day 0, 3 and 4 post infection. Data is the mean of 4 mice per group and error bars represent standard error. C) Kaplan-Meier plot showing percent survival (n=4 mice per group). D) Viral tissue titers in lungs, nasal turbinates, and brains of K18-hACE2 mice harvested at euthanasia (4 dpi for all mice, limit of detection ∼2 log_10_ CCID_50_/g). E) RNA-Seq-derived viral counts per million (CPM) for C57BL/6J and *Ifnar^-/-^* mouse brains 5 days after intranasal infection with SARS-CoV-2_MA1_ (red) or UV-inactivated SARS-CoV-2_MA1_ (blue). Historical data for BA.5 infection in K18-hACE2 mice^30^ is plotted for reference (black circles). There were no statistically significant differences between infectious versus UV-inactivated MA1 for either C56BL/6J or *Ifnar^-/-^* mice. BA.5 versus UV-BA.5 in K18-hACE2 mice was significant by Kolmogorov Smirnov test. The mean and standard error are shown. F) Scatter plots showing principal component 1 (PC1) vs. PC2. Circles represent C57BL/6J mice, and triangles represent *Ifnar^-/-^* mice. Red indicates inoculation with infectious SARS-CoV-2_MA1_, and blue represents inoculation with UV-inactivated SARS-CoV-2_MA1_. G) The log_2_ fold change (x-axis) and –log_10_ FDR (y-axis) are shown for all genes for C57BL/6J mice (top) and *Ifnar^-/-^* mice (bottom). No genes reached the threshold required for statistical significance (dashed line, -log_10_FDR = 1.3), indicating absence of differentially expressed genes (DEGs).

To determine whether SARS-CoV-2_MA1_ increased the propensity for brain infection in a wild-type mouse model, we infected C57BL/6J mice with infectious or UV-inactivated SARS-CoV-2_MA1_. We also infected *Ifnar^-/-^* mice to determine whether neuroinvasiveness of SARS-CoV-2_MA1_ could be enhanced in the absence of competent antiviral type I interferon responses. Mice were humanely euthanized on day 5 post infection as this was expected to be a sufficient timeframe for viral neuroinvasion based on experience in K18-hACE2 mice^30^ (Fig. 3D). Viral titration of lungs and nasal turbinates (Supplementary Figure 2A), and RT-qPCR of lungs (Supplementary Figure 2B), indicated that the respiratory tract of these mice were successfully infected. Some mice contained low viral titers at day 5, consistent with previous experiments in C57BL/6J mice where SARS-CoV-2_MA1_ reached peak viral titers of approximately 10^7^ CCID_50_/g in lungs and nasal turbinates at day 2 post infection, which cleared to undetectable levels in most mice by day 7^9^.

We performed RNA-Seq of C57BL/6J and *Ifnar^-/-^* mouse brains to determine whether there was detectable viral replication or transcriptional responses after infection with SARS-CoV-2_MA1_. UV-inactivated^31,32^ SARS-CoV-2_MA1_ was used as a negative control. There were no significant differences in viral read counts per million (CPM) between mice infected with infectious SARS-CoV-2_MA1_ versus UV-inactivated SARS-CoV-2_MA1_ in either C57BL/6J or*Ifnar^-/-^*mice (Fig. 3E), indicating no detectable virus replication in the brain. In addition, there were less than 1000 viral CPM on average in C57BL/6J and *Ifnar^-/-^* mice, compared to an average of 57000 viral CPM in productively infected K18-hACE2 brains^30^ (Fig. 3E). In addition, there were no separation of treatment groups by principal component analyses (Fig. 3F), and there were no significant DEGs when comparing infectious versus UV-inactivated virus for either mouse strain (Figure 3G, Supplementary Table 3), further supporting absence of viral neuroinvasion and ensuing transcriptional responses. Overall, our data suggests that despite the ability of the spike E484D substitution (as in SARS-CoV-2_MA1_) to facilitate binding to the brain-expressing TMEM106B receptor^13^, there is no overt increased risk of neuroinvasion for SARS-CoV-2_MA1_ in wild-type mice.

### SARS-CoV-2_MA1_ does not replicate in *Ace2^-/-^* mouse lungs

To determine whether ACE2-independent SARS-CoV-2 infection was functional *in vivo*, we infected *Ace2^-/-^* mice (Supplementary Fig. 3 and 4) with SARS-CoV-2_MA1_ and measured viral titers in lungs and nasal turbinates on day 1 and 2 post infection. SARS-CoV-2_MA1_ and SARS-CoV-2_BA.5_ infected C57BL/6J mouse lungs and nasal turbinates, while SARS-CoV-2_QLD02_ (can’t bind mouse ACE2) did not infect C57BL/6J mice (Figure 4A-B). However SARS-CoV-2_MA1_, SARS-CoV-2_BA.5_, or SARS-CoV-2_QLD02_ did not productively infect *Ace2^-/-^* mice (Figure 4A-B, Supplementary Fig. 5). Even when analyzed by the more sensitive RT-qPCR method, there was no detectable SARS-CoV-2_MA1_ replication in *Ace2^-/-^* mice (Figure 4C). SARS-CoV-2_QLD02_ and SARS-CoV-2_BA.5_ had viral RNA levels ∼4 log lower than in C57BL/6J mice, which is likely indicative of background inoculum levels since it is known that these viruses are ACE2-dependent^9^. Overall, these data suggest that SARS-CoV-2 containing the spike E484D substitution do not infect cells via ACE2-independent pathways in mice respiratory tracts.

**Figure 4.**
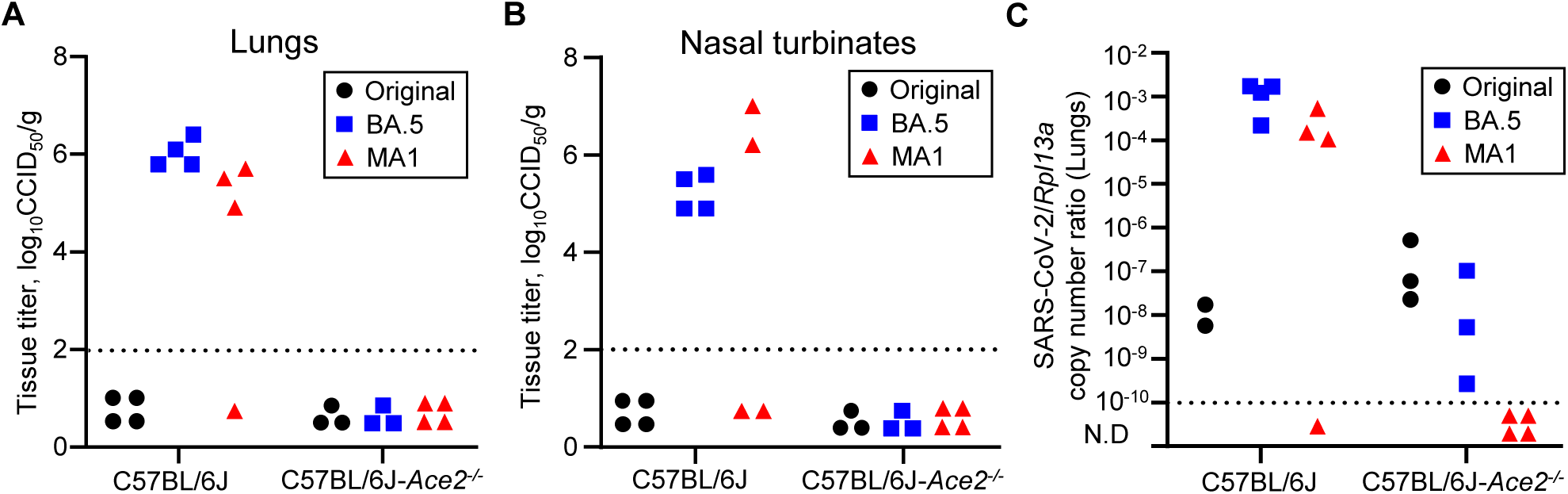
SARS-CoV-2_MA1_ is not capable of ACE2-independent infection in mouse lungs. A) Viral titers in C57BL/6J and *Ace2^-/-^* mouse lungs after 2 days infection with original SARS-CoV-2_QLD02_ (black circles), SARS-CoV-2_BA.5_ (blue squares), or SARS-CoV-2_MA1_ (red triangles). Limit of detection is ∼2 log_10_ CCID_50_/g. B) Viral titers in mouse nasal turbinates after day 2 post infection (same mice as in ‘A’). C) RT-qPCR of RNA from mouse lungs (same mice as in ‘A’) to measure the SARS-CoV-2 RNA copy number levels normalised to the *Rpl13a* housekeeping gene. The dotted line indicates the limit of reliable detection.

## DISCUSSION

Herein we identify that genes involved in heparan sulfate biosynthesis and endocytic pathways were important for ACE2-independent entry of SARS-CoV-2_MA1_, with TMEM106B the major contributor. Baggen *et al*. performed a whole genome CRISPR-Cas9 knockout screen in Huh7 cells using an original SARS-CoV-2 isolate that also identified TMEM106B as important for infection^22^. They showed that Huh7 cells had undetectable ACE2 levels by western blot, and ACE2 was not identified in their screen as important for SARS-CoV-2 infection^22^, together suggesting that their screen was also identifying ACE2-independent infection mechanisms. In a subsequent study, Baggen *et al.* reported that their virus stock acquired the spike E484D substitution after *in vitro* passage during virus stock production^22^, with this substitution a known mediator of ACE2-independent infection *in vitro*^9–12^. Thus, results from our whole genome CRISPR-Cas9 knockout screen which was designed to identify mediators of ACE2-independent infection by SARS-CoV-2_MA1_ is concordant with Baggen *et al*^22^. SARS-CoV-2 with the spike E484D substitution was shown to directly bind TMEM106B^13^. TMEM106B is a type II transmembrane protein predominantly localized in late endosomes and lysosomes with various roles in their function^35^. Small amounts of TMEM106B are expressed on the cell surface which may engage spike-E484D directly as an entry receptor^13^. TMEM106B is also important for post-endocytic fusion, and may also function in spike/TMEM106B mediated cell-cell fusion^13^. Baggen *et al.* also found that heparan sulfates were important for facilitating SARS-CoV-2 entry via either ACE2 or TMEM106B^13^. Overall, our study independently validates the finding that spike E484D substitution allows robust ACE2-independent infection via engaging TMEM106B.

Since TMEM106B is primarily expressed in brain tissue (www.proteinatlas.org) and is involved in neurodegenerative disorders^19,26–29^, we first investigated the prospect that SARS-CoV-2 containing the spike E484D mutation may have increased risk of neurovirulence and/or neuroinvasiveness. Neurovirulence is commonly defined as disease caused by virus replication within the brain, while neuroinvasiveness is defined as virus accessing the brain^36^ which for SARS-CoV-2 is most likely via crossing the cribriform plate and entering the olfactory bulb^33,34,37^. Infection of brain organoids represents a measure of neurovirulence, rather than neuroinvasiveness^30^. SARS-CoV-2_MA1_ replicated in brain organoids significantly higher than natural isolates, although BA.5 and XBB have more productive infection compared to the Original strain^30^. In K18-hACE2 mice, which provides SARS-CoV-2 access to the brain via the olfactory bulb^34^, viral titers in the brain were significantly higher than for the Original strain, perhaps implicating TMEM106B in increased neurovirulence for SARS-CoV-2 strains containing the spike E484D substitution. However interpretation is complicated by K18-hACE2 mice also expressing ACE2 in the brain^38^, thus distinguishing ACE2 from TMEM106B mediated infection would require further investigation. We thus infected C57BL/6J and *Ifnar^-/-^* mice to determine whether SARS-CoV-2_MA1_ could access the brain in mice with endogenous *Ace2* expression, with natural SARS-CoV-2 isolates having low neuroinvasiveness in wild-type mice^30,39^. We previously determined that C57BL/6J mice did not have detectable viral titers in the brain after infection with SARS-CoV-2_MA1_^9^. We now show that even using highly sensitive RNA-Seq, there were no indications that SARS-CoV-2_MA1_ infected the brain or caused transcriptomic changes in the brain of wild-type mice. Others have shown low level SARS-CoV-2 infection in the brains of hamsters, primates and humans^40–43^, suggesting even ACE2-dependent virus can sometimes enter and infect the brain. Overall, our data suggests that SARS-CoV-2_MA1_ may have increased replication within the K18-hACE2 mouse brain, potentially via TMEM106B, but ACE2-independent infection does not provide overt increased opportunities for neuroinvasion.

Prior to our study, ACE2-independent infection has not been demonstrated *in vivo*, thus we generated *Ace2^-/-^* mice on a C57BL/6J background to answer this question. SARS-CoV-2_MA1_ had no detectable replication in *Ace2^-/-^*mouse lungs. Mouse TMEM106B can support ACE2-independent infection^13^, and SARS-CoV-2_MA1_ could infect mouse cell lines^9^, suggesting species differences is unlikely to explain the failure of SARS-CoV-2_MA1_ infection in *Ace2^-/-^* mice. Possible explanations for infection failure include; i) despite TMEM106B mRNA transcripts being detectable in mouse lungs, the protein may not be expressed (as suggested by www.proteinatlas.org), ii) if TMEM106B protein is expressed in mouse lungs, it may not be expressed in the a manner suitable for binding to SARS-CoV-2 spike – for example it may not be expressed on the cell surface, iii) TMEM106B may be expressed in cell types not ordinarily infected by SARS-CoV-2 (e.g. endothelial), while being absent from cell types that are infected by SARS-CoV-2 (e.g. epithelial). Another explanation may be that heparan sulfates are not accessible to SARS-CoV-2 in the respiratory tract, or the interaction of spike with heparan sulfates or TMEM106B are not sufficiently strong to avoid mucocillary clearance before virus entry. The rare studies that have investigated heparan sulfate expression in lungs suggest that it is mainly present on the basal membrane of the epithelium^44,45^, and thus may not be accessible to SARS-CoV-2 spike which binds at the apical epithelium. Our data also hinted at the tenuous nature of ACE2-independent infection of HEK293T cells. While adherent HEK293T cells exhibited robust susceptibility to SARS-CoV-2_MA1_, their vulnerability decreased notably when infection occurred in suspension, irrespective of the detachment method (Supplementary Figure 7). Notably, ACE2-dependent infection remained unaffected by suspension conditions (Supplementary Figure 7). This observation further suggests that the TMEM106B and heparan sulfate entry mechanisms are reliant on adherent cell lines, while ACE2-indepednent entry mechanisms perform poorly in any other setting (e.g. cell suspensions or mouse lungs).

In conclusion, our study identified heparan sulfate and endocytosis, specifically TMEM106B, as key cellular genes/pathways that support ACE2-independent entry by SARS-CoV-2_MA1_ (containing spike E484D substitution). However, ACE2-independent infection by SARS-CoV-2_MA1_ was not supported in lungs of *Ace2^-/-^* mice. Overall, this suggests that robust ACE2-independent infection by SARS-CoV-2_E484D_ is likely an *in vitro* specific phenomenon, and thus emergence of a new variant containing the E484D substitution is unlikely to overtly change the pathogenicity of SARS-CoV-2.

## ACKNOWLEDGEMENTS

The *Ace2* knockout animals were produced via CRISPR genome editing by the Monash Genome Modification Platform (MGMP), Monash University as a node of Phenomics Australia. Phenomics Australia is supported by the Australian Government Department of Education through the National Collaborative Research Infrastructure Strategy, the Super Science Initiative and the Collaborative Research Infrastructure Scheme.

The authors thank the following QIMRB staff; Drs. Anthony White and Lotta Oikari for providing the iPSC line, Dr. Itaru Anraku for management of the PC3 facility at QIMR Berghofer MRI, Dr. Viviana Lutzky for proof reading, Crystal Chang, Blake Ferguson, Ashwini Potadar and Sang-Hee Park for histology services, Tu Parsons and Paul Collins for sequencing services, and the animal house staff for mouse breeding and agistment.

## DATA AVAILABILITY

All data is provided in the manuscript and accompanying supplementary files. Raw sequencing data (fastq files) generated for this publication have been deposited in the NCBI SRA, BioProjects: PRJNA937865 (CRISPR screening), PRJNA911424 (brain organoids), PRJNA1105721 (mouse brains) and are publicly available as of the date of publication.

## AUTHOR CONTRIBUTIONS

Conceptualization, D.J.R.; Methodology, D.J.R.; Formal analysis, D.J.R., T.D. C.B.; Investigation, D.J.R., K.Y., T.D., and B.T.; Resources, D.J.R, and A.S.; Data curation, D.J.R., T.D., and C.B.; Writing, D.J.R. and A.S.; Visualization, D.J.R., C.B., and T.D.; Supervision, D.J.R., Project administration, D.J.R. and A.S.; Funding acquisition, D.J.R., and A.S.

## MATERIALS and METHODS

### Ethics statement and regulatory compliance

All mouse work was conducted in accordance with the “Australian code for the care and use of animals for scientific purposes” as defined by the National Health and Medical Research Council of Australia. Mouse work was approved by the QIMR Berghofer Medical Research Institute Animal Ethics Committee (P3600, A2003-607). For intrapulmonary inoculations, mice were anesthetized using isoflurane. Mice were euthanized using CO_2_.

Breeding and use of GM mice was approved under a Notifiable Low Risk Dealing (NLRD) Identifier: NLRD_Suhrbier_Oct2020: NLRD 1.1(a).

All infectious SARS-CoV-2 work was conducted in a dedicated suite in a biosafety level-3 (PC3) facility at the QIMR Berghofer MRI (Australian Department of Agriculture, Water and the Environment certification Q2326 and Office of the Gene Technology Regulator certification 3445). All work was approved by the QIMR Berghofer MRI Safety Committee (P3600).

### Cell lines, human cortical brain organoids, and virus culture

Vero cells (ATCC#: CCL-81) and HEK293T cells were maintained in RPMI 1640 (Thermo Fisher Scientific, Scoresby, VIC, Australia) and DMEM (Thermo Fisher Scientific, Scoresby, VIC, Australia) supplemented with endotoxin free 10% heat-inactivated fetal bovine serum (FBS; Sigma-Aldrich, Castle Hill, NSW, Australia) at 37°C and 5% CO_2_. Cells were checked for mycoplasma using MycoAlert Mycoplasma Detection Kit (Lonza, Basel, Switzerland).

Human cortical brain organoids (hBOs) were generated and infected exactly as described previously^30^. hBOs were imaged using an EVOS FL (Advanced Microscopy Group), and organoid area within the 2D image circumference was determined by drawing around the edge of the organoid using Image J v1.53^46^.

Virus isolates SARS-CoV-2_QLD02_ (GISAID: EPI_ISL_407896) and SARS-CoV-2_BA.5_ (GISAID: EPI_ISL_15671874, GenBank: OP604184.1), and passage-derived SARS-CoV-2 MA1, were described previously^9,30^. SARS-CoV-2 virus stocks were propagated in Vero E6 (for QLD02 and MA1) or Vero E6-TMPRSS2 (for BA.5) cells^30^. Viruses were titered using CCID_50_ assays on Vero E6 cells^47^.

### Genome wide CRISPR-Cas9 knockout screen

10 x T175 flasks were seeded with 5×10^6^ HEK293T cells per flask overnight. Cells were transduced with the Human CRISPR Knockout Pooled Library^16^ (Brunello, lentiCRISPR v2, Addgene 73179-LV) at ∼MOI 0.3 with 8 µg/ml polybrene. The Brunello CRISPR library contains 74,441 sgRNAs targetting 19,114 protein-coding genes and ≈ 1000 negative control guides, and thus transduction occurred at a rate of approximately 400 cells per sgRNA. After 3 days, transduced cells were selected by adding 1 µg/ml puromycin. After 3-4 days, cells were seeded at 5×10^6^ HEK293T cells per flask overnight. For the uninfected controls, DNA were extracted from 5 flasks using the NucleoSpin Blood Mini kit for DNA from blood as per manufacturers’ instructions (Machery Nagel). Another 16 flasks were infected with SARS-CoV-2_MA1_ at MOI 0.1 and incubated for 3 days at 37°C and 5% CO_2_. Cytopathic effect was evident, with very few cells remaining attached to the flask. Media was removed and discarded and flasks gently washed using PBS. Cells were maintained for 21 days to allow recovery and expansion of surviving cells, and DNA was purified from cells using the NucleoSpin Blood Mini kit for DNA from blood as per manufacturers’ instructions (Machery Nagel). Cells were also seeded and re-infected with SARS-CoV-2_MA1_, with no overt cytopathic effect observed (suggesting resistance to SARS-CoV-2_MA1_ infection).

The sgRNA-containing region of the lentivirus in the sample DNA was PCR amplified using NEBNext Ultra II Q5 Master Mix as per manufacturers’ instructions (New England Biolabs), and with primers (Integrated DNA Technologies) containing Illumina adapters and TruSeq indices according to the Broad Institute protocol “PCR of sgRNAs for Illumina Sequencing”. PCR products were purified using AMPure XP Beads as per manufacturers’ instructions (Beckman Coulter). PCR products were sequenced using the llumina NextSeq 550 platform with 5% PhiX (Illumina) spike-in, generating 75 bp single end reads.

For bioinformatic data analyses, the Robust Rank Aggregation algorithm in the MAGeCK software^48^ was used with default settings to rank the sgRNAs that are significantly enriched or depleted (q<0.05, FDR by Benjamini-Hochberg method) (see Supplementary Table 1).

### Growth kinetics in RENcell VM

RENCell VM were cultured as described previously^49^. Briefly, cells were seeded at 50,000 cells per well in 12 well plates by detaching cells using StemPro Accutase Cell Dissociation Reagent (Thermo Fisher Scientific), and seeding in plates coated with Matrigel (Sigma). RENcells were infected with MA1, BA1 and BA5 at multiplicity of infection (MOI) 1, and after one hour virus was removed and cells were washed in PBS once before replacing culture media (DMEM F-12 (Thermo Fisher Scientific), penicillin (100LIU/ml)/streptomycin (100Lμg/ml) (Gibco/Life Technologies), 20 ng/ml FGF (STEMCELL Technologies), 20 ng/ml EGF (STEMCELL Technologies), and 2% B27 supplements (Thermo Fisher Scientific). Supernatant was collected daily and virus titer determined by CCID_50_ assays. For crystal violet staining of remaining cells after infection, formaldehyde (7.5% w/v)/crystal violet (0.05% w/v) was added to wells overnight, plates washed twice in water, and plates dried overnight.

### Mouse infection and monitoring

Female C57BL/6J (10-week old to 50-week old) mice were purchased from Ozgene (Perth, Western Australia, Australia). K18-hACE2 mice (strain B6.Cg-Tg(K18-ACE2)2Prlmn/J, JAX Stock No: 034860) were purchased from The Jackson Laboratory, and were bred to homozygotes as described previously^30^. *Ifnar^-/-^* mice (originally provided by P. Hertzog, Monash University, Melbourne, VIC, Australia^50^) on a C57BL/6J background were bred in-house^51^. The Australian Phenomics Network, Monash University, Melbourne, Australia, used CRISPR to generate *Ace2^+/-^* mice as described in Supplementary Figure 3, which were subsequently bred to homozygotes (Supplementary Figure 4) and maintained in house at QIMRB. The *Ace2^-/-^* mouse line was donated to The Jackson Laboratory, and are available under the strain number 038377. All mice at QIMRB were held under standard animal house conditions (for details see^9^).

For each experiment, mice were sorted into groups with a similar age distribution. Mice were infected intrapulmonary via the nasal route with 5×10^4^ CCID_50_ of virus in 50 μl RPMI 1640 medium while under light anaesthesia; 3 % isoflurane (Piramal Enterprises Ltd., Andhra Pradesh, India) delivered using TheStinger, Rodent Anaesthesia System (Advanced Anaesthesia Specialists/Darvall, Gladesville, NSW, Australia). Mice were weighed and overt disease symptoms scored as described^33^. Mice were euthanized using CO2, and tissues (lungs, brains, nasal turbinates) were homogenized using four ceramic beads at 6000 rpm twice for 15s (Precellys 24 Homogenizer, Bertin Instruments, Montigny-le-Bretonneux, France). After centrifugation for 10 min, 9400 ×g at 4°C, virus titers in supernatants were determined by CCID_50_ assays using Vero E6 cells. Where applicable, tissues were placed in RNAlater (Sigma Aldrich) at 4°C overnight then stored at -80°C for RNA extraction and sequencing, or 10% formalin for histology.

### Histology and immunohistochemistry

Mouse lungs and brain organoids were fixed in 10 % formalin, embedded in paraffin, and sections stained with H&E (Sigma-Aldrich, Darmstadt, Germany). Slides were scanned using Aperio AT Turbo (Aperio, Vista, CA, USA) and images extracted using Aperio Image Scope software v12.3.2.8013 (Leica Biosystems, Wetzlar, Germany). Immunohistochemistry was undertaken as described using the in-house developed anti-spike monoclonal antibody 1E8^30,31^, or an ACE2 antibody (Abcam, ab108209, EPR4436)^32^.

### RT-qPCR of mouse lungs

Mouse lungs were transferred from RNAlater to TRIzol (Life Technologies) and werehomogenized twice at 6000 x g for 15 sec. Homogenates were centrifuged at 14,000 × g for 10 min and RNA was isolated as per manufacturers’ instructions. cDNA was synthesized using ProtoScript II First Strand cDNA Synthesis Kit (New England Biolabs) and qPCR performed using iTaq Universal Probes Kit (Bio-Rad) for SARS-CoV-2 primers^52^, or iTaq Universal SYBR Green Supermix (Bio-Rad) for mRPL13a primers. The following primer/probes (Integrated DNA Technologies Australia, Sydney, Australia) were used; SARS-CoV-2 (Sarbeco-E^53^) Forward 5′-ACAGGTACGTTAATAGTTAATAGCGT-3′, Reverse 5′-ATATTGCAGCAGTACG CACACA -3’, and Probe 5’-FAM-ACACTAGCCATCCTTACTGCGCTTCG-ZEN™/Iowa Black® FQ -3’. mRPL13a Forward 5′-GAGGTCGGGTGGAAGTACCA-3′, Reverse 5′-TGCATCTTGGCCTTTTCCTT-3′. Reactions were placed into a BioRad CFX96 based on the following cycling protocol: 10 min at 50°C, 3 min at 95°C, and 40 cycles of 15 s at 95°C and 30 s at 60°C. Copy number was calculated from a standard curve of purified PCR products or plasmids and normalised to mRPL13a as described^32^.

### RNA-Seq and bioinformatics

Total RNA was extracted from mouse brains and human brain organoids using TRIzol. RNA concentration and quality was measured using TapeStation D1K TapeScreen assay (Agilent). cDNA libraries were prepared using the Illumina TruSeq Stranded mRNA library prep kit, and the sequencing performed on the Illumina Nextseq 550 platform generating 75 bp paired end reads. The quality of raw sequencing reads was assessed using FastQC v0.11.8 and trimmed using Cutadapt v2.3 to remove adapter sequences and low-quality bases. Trimmed reads were aligned using STAR v2.7.1a to a combined reference that included the host (GRCm38 primary assembly for mouse brain samples and GRCh38 for human brain organoid samples) and the SARS-CoV-2 Wuhan (NC_045512.2) reference genome. Gene expression was estimated using RSEM v1.3.0. Reads aligned to SARS-Cov-2 were counted using SAMtools v1.9. Differential gene expression was analyzed using EdgeR v3.42.4 in R v4.1.0. Significantly differentially expressed genes (DEGs) were identified using a FDR cutoff of <0.05. Principle components analyses of log2 normalised counts was performed using R and ggplot2. Pathway analysis was performed on DEGs using Ingenuity Pathway Analysis v111725566 (QIAGEN).

### Statistics

Statistical analyses of experimental data were performed using IBM SPSS Statistics for Windows, Version 19.0 (IBM Corp., Armonk, NY, USA). The t-test was used when the difference in variances was <4, skewness was > -2 and kurtosis was <2. Otherwise, the non-parametric Kolmogorov-Smirnov test was used.

---

**Supplementary Figure 1.**
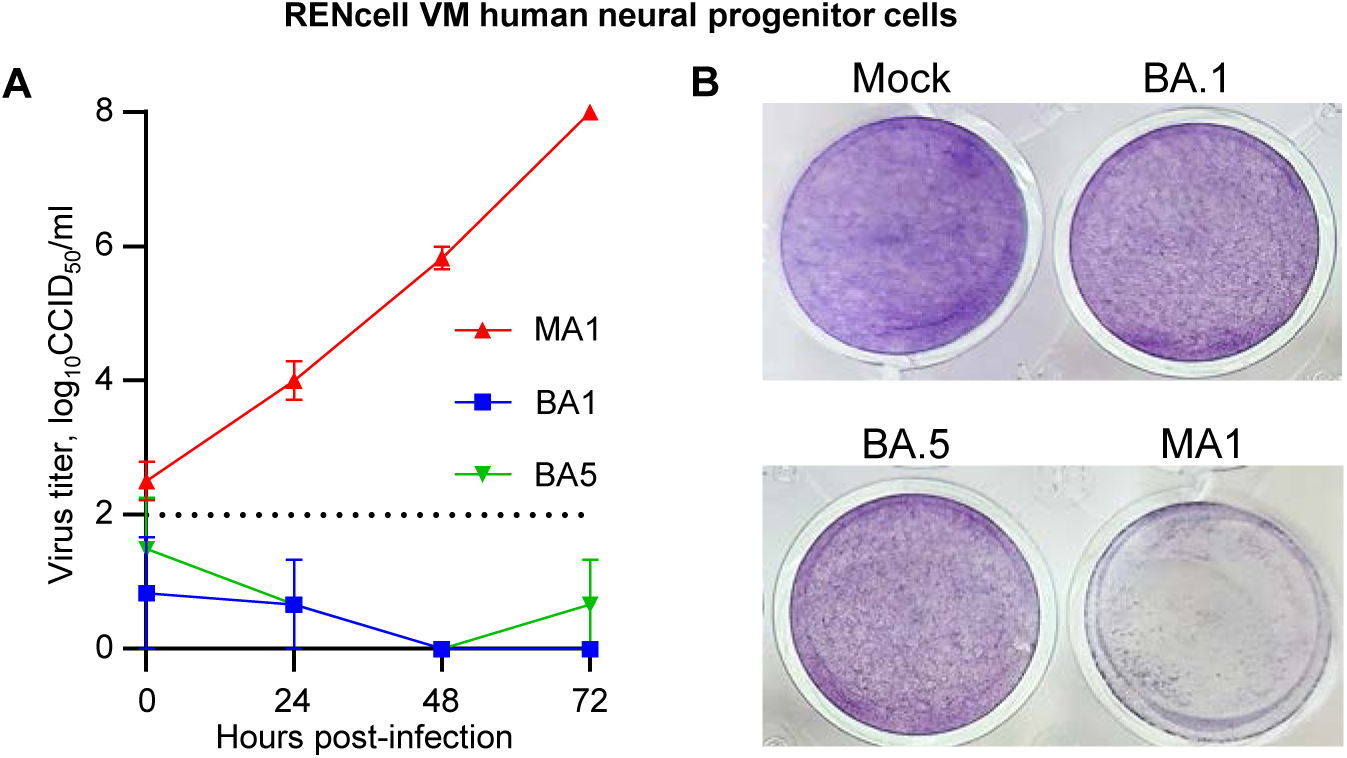
Replication and CPE in RENcell VM neural progenitor cells. **A)** Viral growth kinetics in RENcell VM determined by CCID_50_ assays of culture supernatant at the indicated times post infection. Data is the mean of 3 replicates per virus isolate; (limit of detection is 2 log_10_CCID_50_/ml). **B)** Images of crystal violet stained RENcell VM at 6 dpi (representative of n=3 per group). Less blue/violet stained cells indicates more viral CPE.

**Supplementary Fig. 2.**
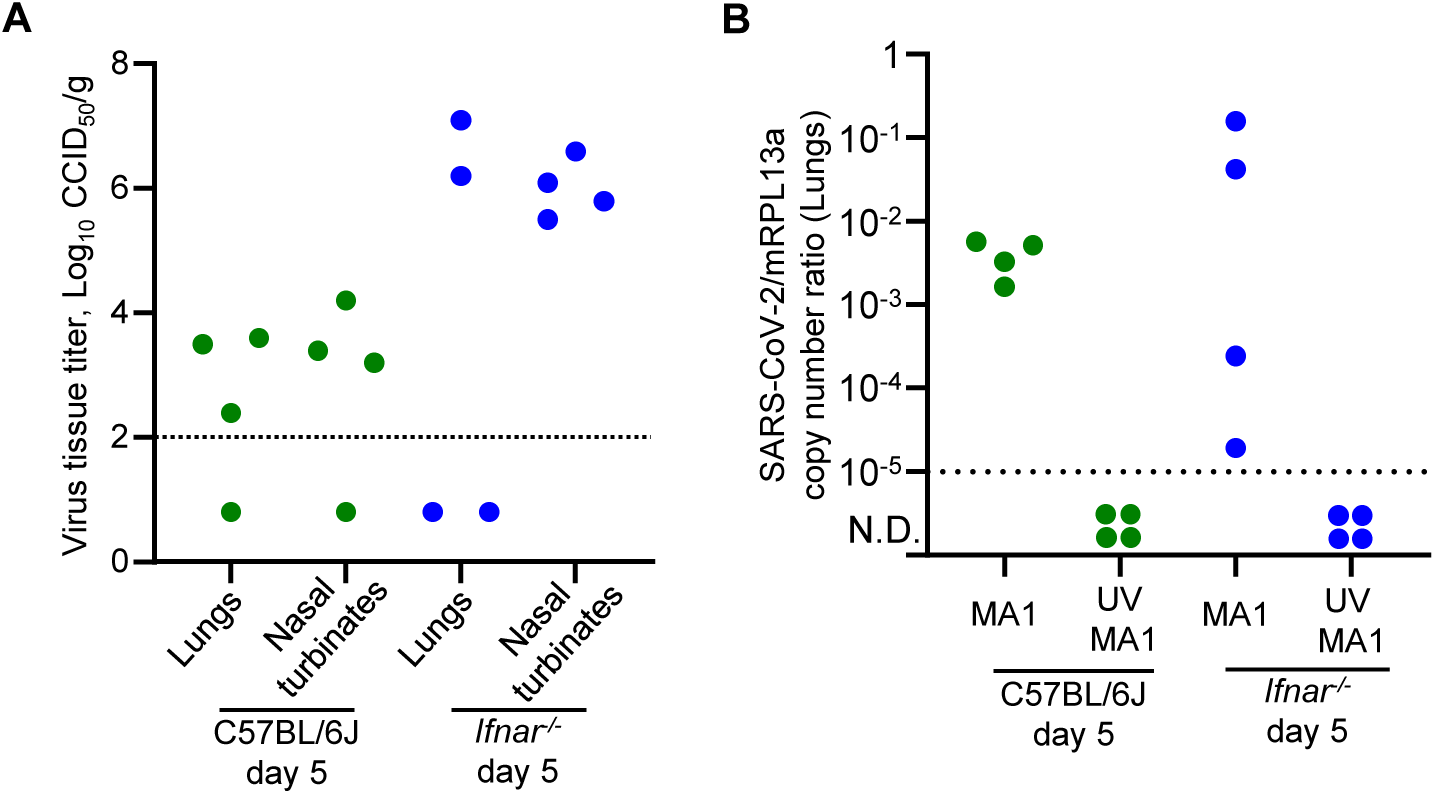
Confirmation of successful lung and nasal turbinate infection in C57BL/6J and *Ifnar^-/-^* mice. Mouse lungs and nasal turbinates were collected on day 5 for viral titrations, and mouse lungs were also collected for RT-qPCR, from the same mice shown in Figure 3E (brain RNA-Seq). **A)** Viral titers in mouse lungs and nasal turbinates were determined by CCID_50_ assay. Viral titers for some mice fell below the limit of detection (2 log_10_ CCID_50_/ml), indicating infectious virus had been cleared in some mice by day 5. **B)** RT-qPCR is a more sensitive method at detecting viral infection in lungs. All SARS-CoV-2_MA1_ infected mouse lungs had detectable viral RNA, while UV-inactivated virus inoculated mice all had undetectable lung titers. This indicates that all mice that received a SARS-CoV-2_MA1_ inoculum established a successful lung infection.

**Supplementary Fig. 3.**
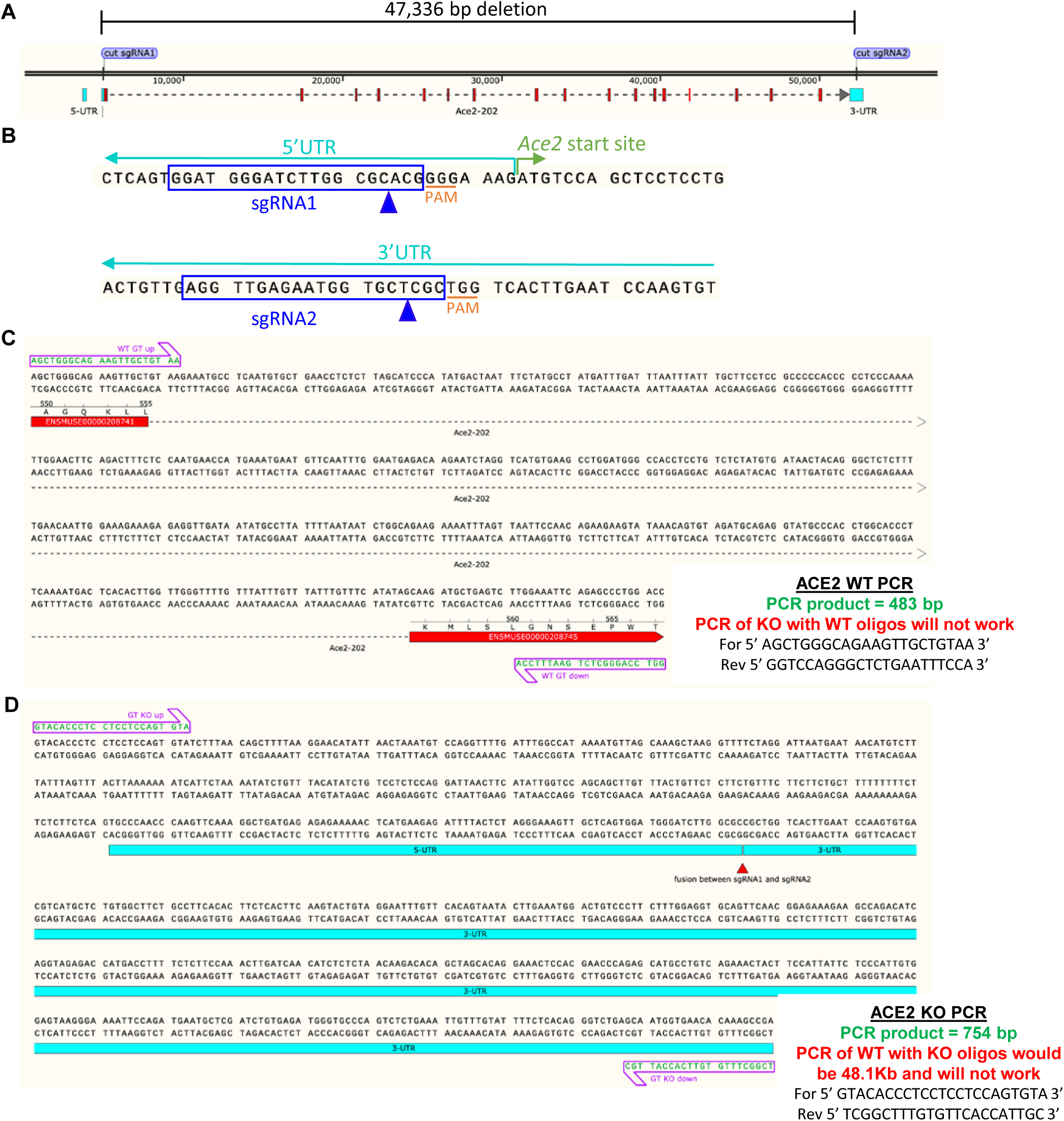
Generation of *Ace2^-/-^* mice. **A)** Schematic showing deletion of all *Ace2* exons. **B)** sgRNAs and location of cut sites (blue arrow heads). **C)** Primers for genotyping WT allele. **D)** Primers for genotyping *Ace2^-/-^* allele.

**Supplementary Fig. 4.**
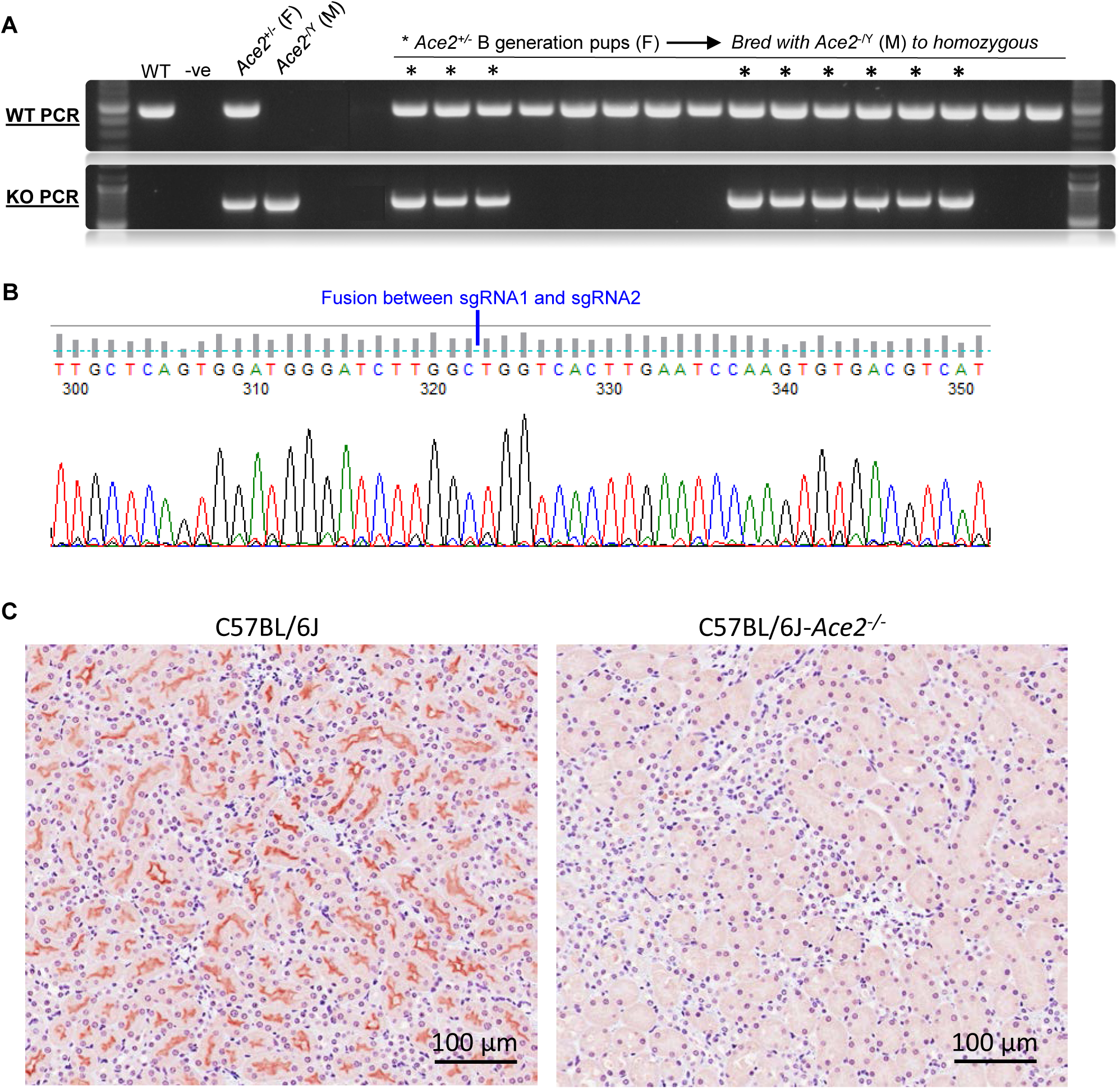
Validation of *Ace2^-/-^* mice. **A)** Example of WT or KO PCR products for second generation *Ace2^+/-^* pups. The *Ace2^-/Y^* male was bred with *Ace2^+/-^* generation B females to generate a stable homozygous *Ace2^-/-^* line. **B)** Example of sequencing 754 bp KO PCR product, showing the fusion site after sgRNA1 and sgRNA2 guided deletion of the *Ace2* coding sequence. **C)** Immunohistochemistry of kidneys from C57BL/6J or C57BL/6J-*Ace2^-/-^* mice using anti-Ace2 antibody (Abcam, ab108209). Brown staining indicates Ace2 expression in C57BL/6J mice, but not in C57BL/6J-*Ace2^-/-^* mice.

**Supplementary Fig. 5.**
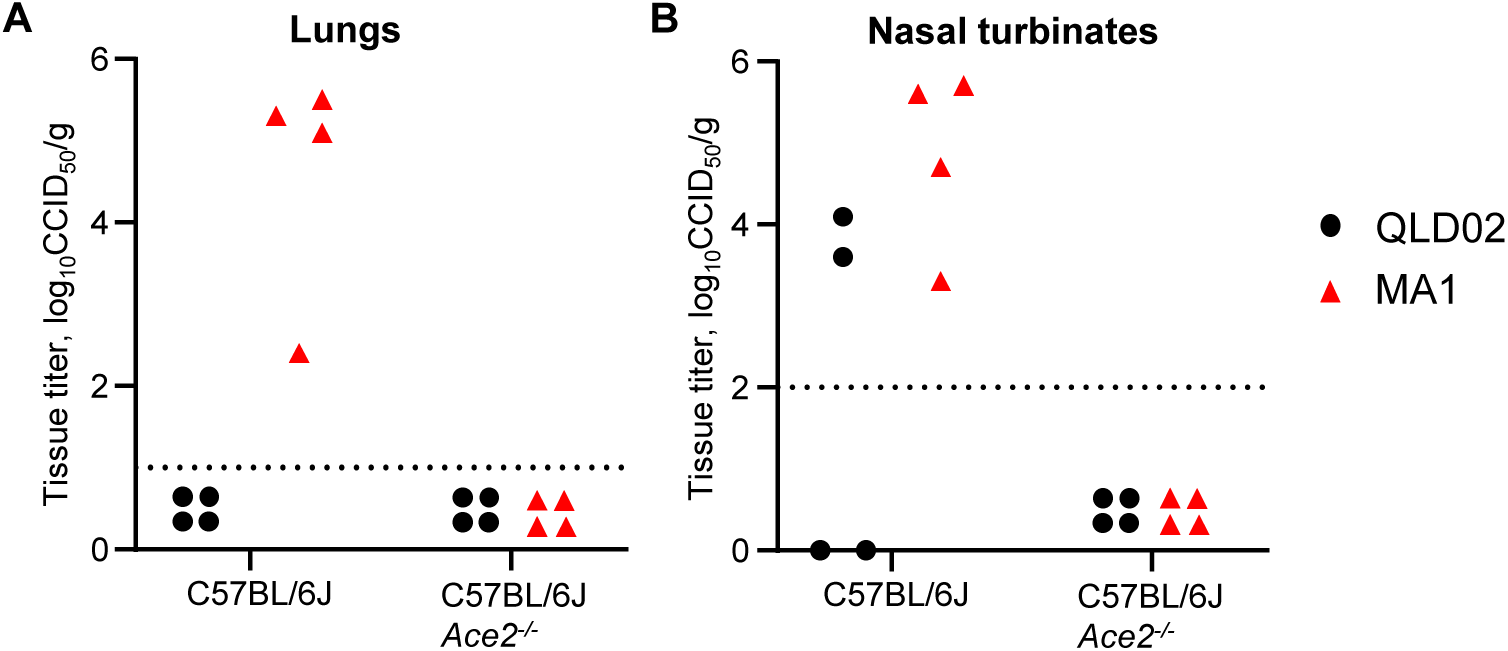
Day 1 post-infection timepoint for *Ace2^-/-^* mice. C57BL/6J mice or C57BL/6J-*Ace2^-/-^* mice were infected with SARS-CoV-2_QLD02_ or SARS-CoV-2_MA1_, and **(A)** lungs or **(B)** nasal turbinates were collected at 1 day post-infection for virus titrations by CCID_50_ assay.

**Supplementary Fig. 6.**
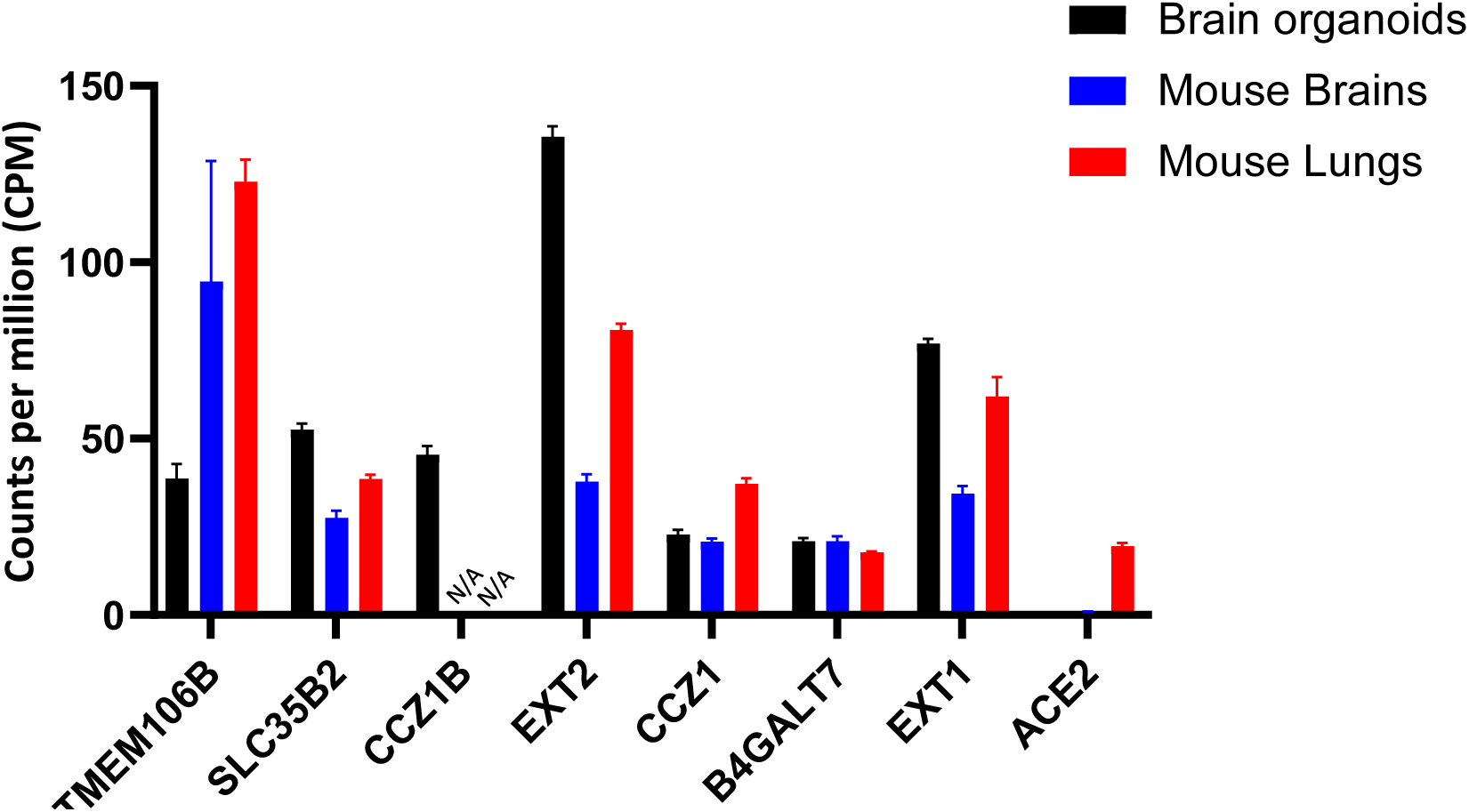
Gene expression by RNA-Seq for CRISPR screening hits in brain organoids, mouse brains, and mouse lungs.

**Supplementary Fig. 7.**
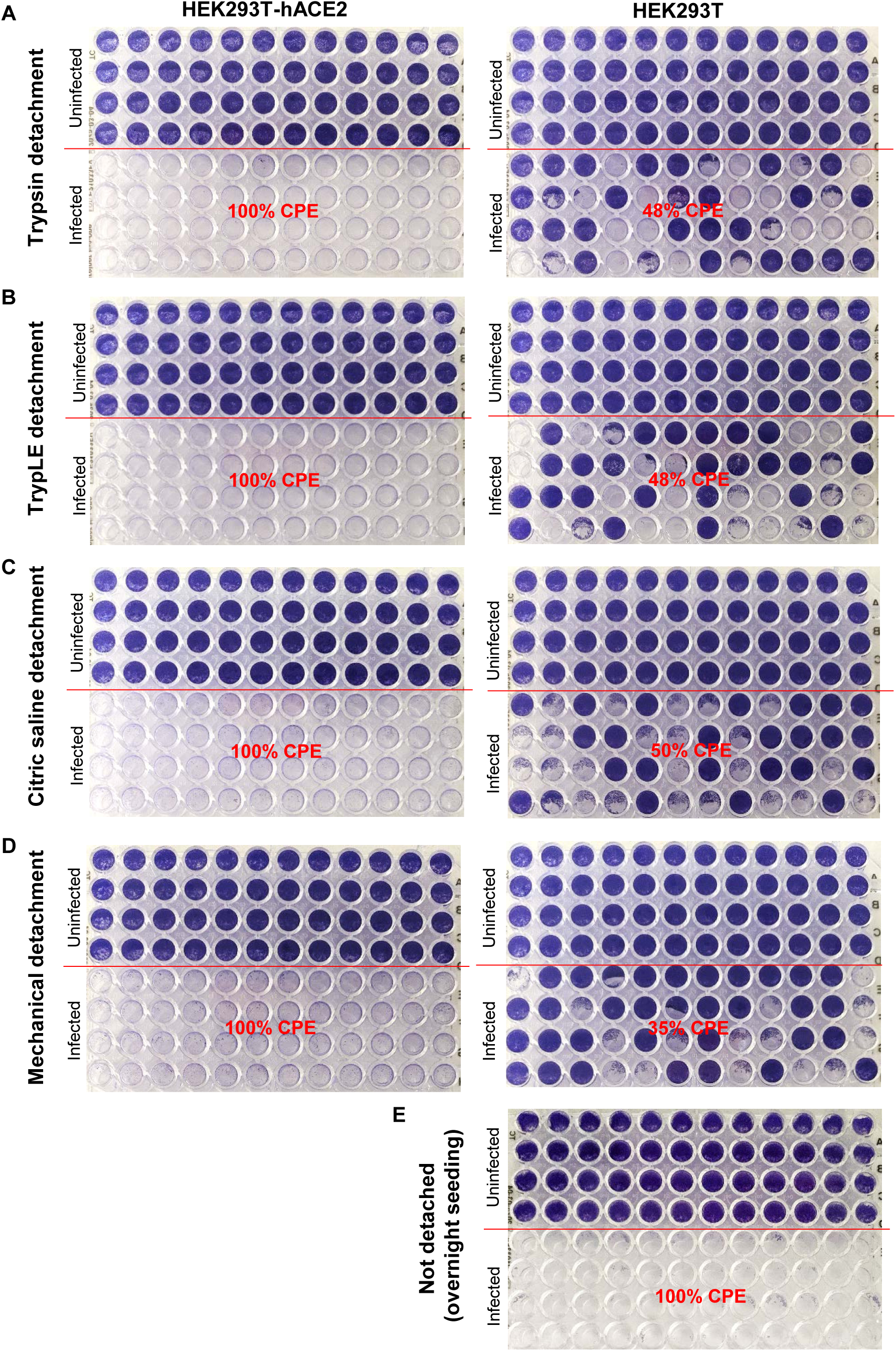
ACE-independent infection is less efficient when cells are infected in suspension. HEK293T-hACE2 (left) or HEK293T (right) cells (10,000 per well of 96 well plates) were infected with 100 CCID_50_ SARS-CoV-2_MA1_ either in suspension **(A-D)**, or after seeding overnight **(E)**. Cells in suspension were detached immediately prior to infection using **(A)** trypsin, **(B)** TrpLE (Thermo Fisher Scientific), **(C)** citric saline, or **(D)** mechanical detachment by gently resuspending in media. After 4 days, plates were fixed and stained with crystal violet to show cytopathic effect (CPE) caused by the virus infection.

## REFERENCES

1 Hoffmann, M. et al. SARS-CoV-2 Cell Entry Depends on ACE2 and TMPRSS2 and Is Blocked by a Clinically Proven Protease Inhibitor. Cell 181, 271–280.e278, 10.1016/j.cell.2020.02.052 (2020).

2 Jackson, C. B., Farzan, M., Chen, B. & Choe, H. Mechanisms of SARS-CoV-2 entry into cells. Nature Reviews Molecular Cell Biology 23, 3–20, doi:10.1038/s41580-021-00418-x (2022).

3 Peacock, T. P. et al. The furin cleavage site in the SARS-CoV-2 spike protein is required for transmission in ferrets. Nature Microbiology 6, 899–909, doi:10.1038/s41564-021-00908-w (2021).

4 Lim, S., Zhang, M. & Chang, T. L. ACE2-Independent Alternative Receptors for SARS-CoV-2. Viruses 14 (2022).

5 Amraei, R. et al. CD209L/L-SIGN and CD209/DC-SIGN Act as Receptors for SARS-CoV-2. ACS Cent Sci 7, 1156–1165, doi:10.1021/acscentsci.0c01537 (2021).

6 Wang, S. et al. AXL is a candidate receptor for SARS-CoV-2 that promotes infection of pulmonary and bronchial epithelial cells. Cell Research 31, 126–140, doi:10.1038/s41422-020-00460-y (2021).

7 Cantuti-Castelvetri, L. et al. Neuropilin-1 facilitates SARS-CoV-2 cell entry and infectivity. Science 370, 856–860, doi:10.1126/science.abd2985 (2020).

8 Wang, K. et al. CD147-spike protein is a novel route for SARS-CoV-2 infection to host cells. Signal Transduction and Targeted Therapy 5, 283, doi:10.1038/s41392-020-00426-x (2020).

9 Yan, K. et al. Evolution of ACE2-independent SARS-CoV-2 infection and mouse adaption after passage in cells expressing human and mouse ACE2. Virus Evolution 8, veac063, doi:10.1093/ve/veac063 (2022).

10 Ramirez, S. et al. Overcoming Culture Restriction for SARS-CoV-2 in Human Cells Facilitates the Screening of Compounds Inhibiting Viral Replication. Antimicrobial Agents and Chemotherapy 65, 10.1128/aac.00097-00021, doi:10.1128/aac.00097-21 (2021).

11 Hoffmann, M. et al. Evidence for an ACE2-Independent Entry Pathway That Can Protect from Neutralization by an Antibody Used for COVID-19 Therapy. mBio 13, e0036422, doi:10.1128/mbio.00364-22 (2022).

12 Puray-Chavez, M. et al. Systematic analysis of SARS-CoV-2 infection of an ACE2-negative human airway cell. Cell Rep 36, 109364, doi:10.1016/j.celrep.2021.109364 (2021).

13 Baggen, J. et al. TMEM106B is a receptor mediating ACE2-independent SARS-CoV-2 cell entry. Cell 186, 3427–3442.e3422, 10.1016/j.cell.2023.06.005 (2023).

14 Vu, M. N. et al. QTQTN motif upstream of the furin-cleavage site plays a key role in SARS-CoV-2 infection and pathogenesis. Proceedings of the National Academy of Sciences 119, e2205690119, doi:10.1073/pnas.2205690119 (2022).

15 Liu, Z. et al. Identification of Common Deletions in the Spike Protein of Severe Acute Respiratory Syndrome Coronavirus 2. J Virol 94, doi:10.1128/jvi.00790-20 (2020).

16 Doench, J. G. et al. Optimized sgRNA design to maximize activity and minimize off-target effects of CRISPR-Cas9. Nat Biotechnol 34, 184–191, doi:10.1038/nbt.3437 (2016).

17 McCormick, C., Duncan, G., Goutsos, K. T. & Tufaro, F. The putative tumor suppressors EXT1 and EXT2 form a stable complex that accumulates in the Golgi apparatus and catalyzes the synthesis of heparan sulfate. Proceedings of the National Academy of Sciences 97, 668–673, doi:10.1073/pnas.97.2.668 (2000).

18 Monteil, V. et al. Identification of CCZ1 as an essential lysosomal trafficking regulator in Marburg and Ebola virus infections. Nature Communications 14, 6785, doi:10.1038/s41467-023-42526-6 (2023).

19 Jiao, H.-S., Yuan, P. & Yu, J.-T. TMEM106B aggregation in neurodegenerative diseases: linking genetics to function. Molecular Neurodegeneration 18, 54, doi:10.1186/s13024-023-00644-1 (2023).

20 Rebendenne, A. et al. Bidirectional genome-wide CRISPR screens reveal host factors regulating SARS-CoV-2, MERS-CoV and seasonal HCoVs. Nature Genetics 54, 1090–1102, doi:10.1038/s41588-022-01110-2 (2022).

21 Grodzki, M. et al. Genome-scale CRISPR screens identify host factors that promote human coronavirus infection. Genome Medicine 14, 10, doi:10.1186/s13073-022-01013-1 (2022).

22 Baggen, J. et al. Genome-wide CRISPR screening identifies TMEM106B as a proviral host factor for SARS-CoV-2. Nature Genetics 53, 435–444, doi:10.1038/s41588-021-00805-2 (2021).

23 Zhu, Y. et al. A genome-wide CRISPR screen identifies host factors that regulate SARS-CoV-2 entry. Nature Communications 12, 961, doi:10.1038/s41467-021-21213-4 (2021).

24 Synowiec, A. et al. Identification of Cellular Factors Required for SARS-CoV-2 Replication. Cells 10, doi:10.3390/cells10113159 (2021).

25 Wang, R. et al. Genetic Screens Identify Host Factors for SARS-CoV-2 and Common Cold Coronaviruses. Cell 184, 106–119.e114, 10.1016/j.cell.2020.12.004 (2021).

26 Werner, G. et al. Loss of TMEM106B potentiates lysosomal and FTLD-like pathology in progranulin-deficient mice. EMBO Rep 21, e50241, doi:10.15252/embr.202050241 (2020).

27 Brady, O. A., Zheng, Y., Murphy, K., Huang, M. & Hu, F. The frontotemporal lobar degeneration risk factor, TMEM106B, regulates lysosomal morphology and function. Hum Mol Genet 22, 685–695, doi:10.1093/hmg/dds475 (2013).

28 Zhang, T. et al. TMEM106B regulates microglial proliferation and survival in response to demyelination. Science Advances 9, eadd2676, doi:10.1126/sciadv.add2676 (2023).

29 Simons, C. et al. A recurrent de novo mutation in TMEM106B causes hypomyelinating leukodystrophy. Brain 140, 3105–3111, doi:10.1093/brain/awx314 (2017).

30 Stewart, R. et al. SARS-CoV-2 omicron BA.5 and XBB variants have increased neurotropic potential over BA.1 in K18-hACE2 mice and human brain organoids. Frontiers in Microbiology 14, doi:10.3389/fmicb.2023.1320856 (2023).

31 Morgan, M. S. et al. Monoclonal Antibodies Specific for SARS-CoV-2 Spike Protein Suitable for Multiple Applications for Current Variants of Concern. Viruses 15, doi:10.3390/v15010139 (2022).

32 Rawle, D. J. et al. ACE2-lentiviral transduction enables mouse SARS-CoV-2 infection and mapping of receptor interactions. PLoS Pathog 17, e1009723, doi:10.1371/journal.ppat.1009723 (2021).

33 Dumenil, T. et al. Warmer ambient air temperatures reduce nasal turbinate and brain infection, but increase lung inflammation in the K18-hACE2 mouse model of COVID-19. Sci Total Environ 859, 160163, doi:10.1016/j.scitotenv.2022.160163 (2023).

34 Fumagalli, V. et al. Administration of aerosolized SARS-CoV-2 to K18-hACE2 mice uncouples respiratory infection from fatal neuroinvasion. Sci Immunol 7, eabl9929, doi:10.1126/sciimmunol.abl9929 (2022).

35 Klein, Z. A. et al. Loss of TMEM106B Ameliorates Lysosomal and Frontotemporal Dementia-Related Phenotypes in Progranulin-Deficient Mice. Neuron 95, 281–296.e286, doi:10.1016/j.neuron.2017.06.026 (2017).

36 Bauer, L. et al. The neuroinvasiveness, neurotropism, and neurovirulence of SARS-CoV-2. Trends in Neurosciences 45, 358–368, 10.1016/j.tins.2022.02.006 (2022).

37 Jiao, L. et al. The olfactory route is a potential way for SARS-CoV-2 to invade the central nervous system of rhesus monkeys. Signal Transduct Target Ther 6, 169, doi:10.1038/s41392-021-00591-7 (2021).

38 Carossino, M. et al. Fatal Neurodissemination and SARS-CoV-2 Tropism in K18-hACE2 Mice Is Only Partially Dependent on hACE2 Expression. Viruses 14, doi:10.3390/v14030535 (2022).

39 Pan, T. et al. Infection of wild-type mice by SARS-CoV-2 B.1.351 variant indicates a possible novel cross-species transmission route. Signal Transduction and Targeted Therapy 6, 420, doi:10.1038/s41392-021-00848-1 (2021).

40 Stein, S. R. et al. SARS-CoV-2 infection and persistence in the human body and brain at autopsy. Nature 612, 758–763, doi:10.1038/s41586-022-05542-y (2022).

41 Beckman, D. et al. SARS-CoV-2 infects neurons and induces neuroinflammation in a non-human primate model of COVID-19. Cell Rep 41, 111573, doi:10.1016/j.celrep.2022.111573 (2022).

42 de Melo, G. D. et al. Neuroinvasion and anosmia are independent phenomena upon infection with SARS-CoV-2 and its variants. Nat Commun 14, 4485, doi:10.1038/s41467-023-40228-7 (2023).

43 Proal, A. D. et al. SARS-CoV-2 reservoir in post-acute sequelae of COVID-19 (PASC). Nature Immunology 24, 1616–1627, doi:10.1038/s41590-023-01601-2 (2023).

44 He, H., Huang, M., Sun, S., Wu, Y. & Lin, X. Epithelial heparan sulfate regulates Sonic Hedgehog signaling in lung development. PLOS Genetics 13, e1006992, doi:10.1371/journal.pgen.1006992 (2017).

45 Gill, S., Wight, T. N. & Frevert, C. W. Proteoglycans: key regulators of pulmonary inflammation and the innate immune response to lung infection. Anat Rec (Hoboken) 293, 968–981, doi:10.1002/ar.21094 (2010).

46 Schneider, C. A., Rasband, W. S. & Eliceiri, K. W. NIH Image to ImageJ: 25 years of image analysis. Nature Methods 9, 671–675, doi:10.1038/nmeth.2089 (2012).

47 Yan, K., Rawle, D. J., Le, T. T. & Suhrbier, A. Simple rapid in vitro screening method for SARS-CoV-2 anti-virals that identifies potential cytomorbidity-associated false positives. Virol J 18, 123, doi:10.1186/s12985-021-01587-z (2021).

48 Li, W. et al. MAGeCK enables robust identification of essential genes from genome-scale CRISPR/Cas9 knockout screens. Genome Biol 15, 554, doi:10.1186/s13059-014-0554-4 (2014).

49 Wilson, N. et al. Neurovirulence of the Australian outbreak Japanese Encephalitis virus genotype 4 is lower compared to genotypes 2 and 3 in mice and human cortical brain organoids. bioRxiv, 2023.2004.2026.538504, doi:10.1101/2023.04.26.538504 (2023).

50 Swann, J. B. et al. Type I IFN contributes to NK cell homeostasis, activation, and antitumor function. J Immunol 178, 7540–7549, doi:10.4049/jimmunol.178.12.7540 (2007).

51 Hazlewood, J. E. et al. The Chimeric Binjari-Zika Vaccine Provides Long-Term Protection against ZIKA Virus Challenge. Vaccines (Basel) 10, doi:10.3390/vaccines10010085 (2022).

52 Pollak, N. M. et al. Rapid inactivation and sample preparation for SARS-CoV-2 PCR-based diagnostics using TNA-Cifer Reagent E. Frontiers in Microbiology 14, doi:10.3389/fmicb.2023.1238542 (2023).

53 Vogels, C. B. F. et al. Analytical sensitivity and efficiency comparisons of SARS-CoV-2 RT-qPCR primer-probe sets. Nat Microbiol 5, 1299–1305, doi:10.1038/s41564-020-0761-6 (2020).

